# Automated ChIPmentation procedure on limited biological material of the human blood fluke Schistosoma mansoni

**DOI:** 10.1101/2021.06.02.446743

**Authors:** Chrystelle Lasica, Ronaldo de Carvalho Augusto, Helene Mone, Gabriel Mouahid, Cristian Chaparro, Anne-Clemence Veillard, Agnieszka Zelisko-Schmidt, Christoph Grunau

## Abstract

Automated ChIPmentation procedure is a convenient alternative to native chromatin immunoprecipitation (N-ChIP). It is now routinely used for ChIP-Seq. Using the human parasite *Schistosoma mansoni*, whose production requires scarifying animals and should therefore kept to a minimum, we show here that the automated ChIPmentation is suitable for limited biological material. We define as operational limit ≥20,000 cells. We also present a streamlined protocol for the preparation of ChIP input libraries.

## Introduction

*Schistosoma mansoni* is a human parasite with a complex life cycle that shows strong developmental phenotypic plasticity, with intra-molluscal, and intra-vertebrate stages and two free-swimming larvae stages (miracidium and cercariae). We had shown by native chromatin immunoprecipitation (N-ChIP) that the different life cycle stages show also strong histone modification plasticity (Cosseau et al. 2009, Roquis et al. 2018, de Carvalho Augusto et al. 2020). While N-ChIP is in principle doable we found that there are two challenges associated with: one is the high hands-on time with the N-ChIP and the other is to obtain enough biological material for performing several ChIP experiments with different antibodies. We therefore explored an automated ChIP procedure (Figure 1) and we tested what the lowest reasonable cell number would be with this procedure.

**Figure 1:**
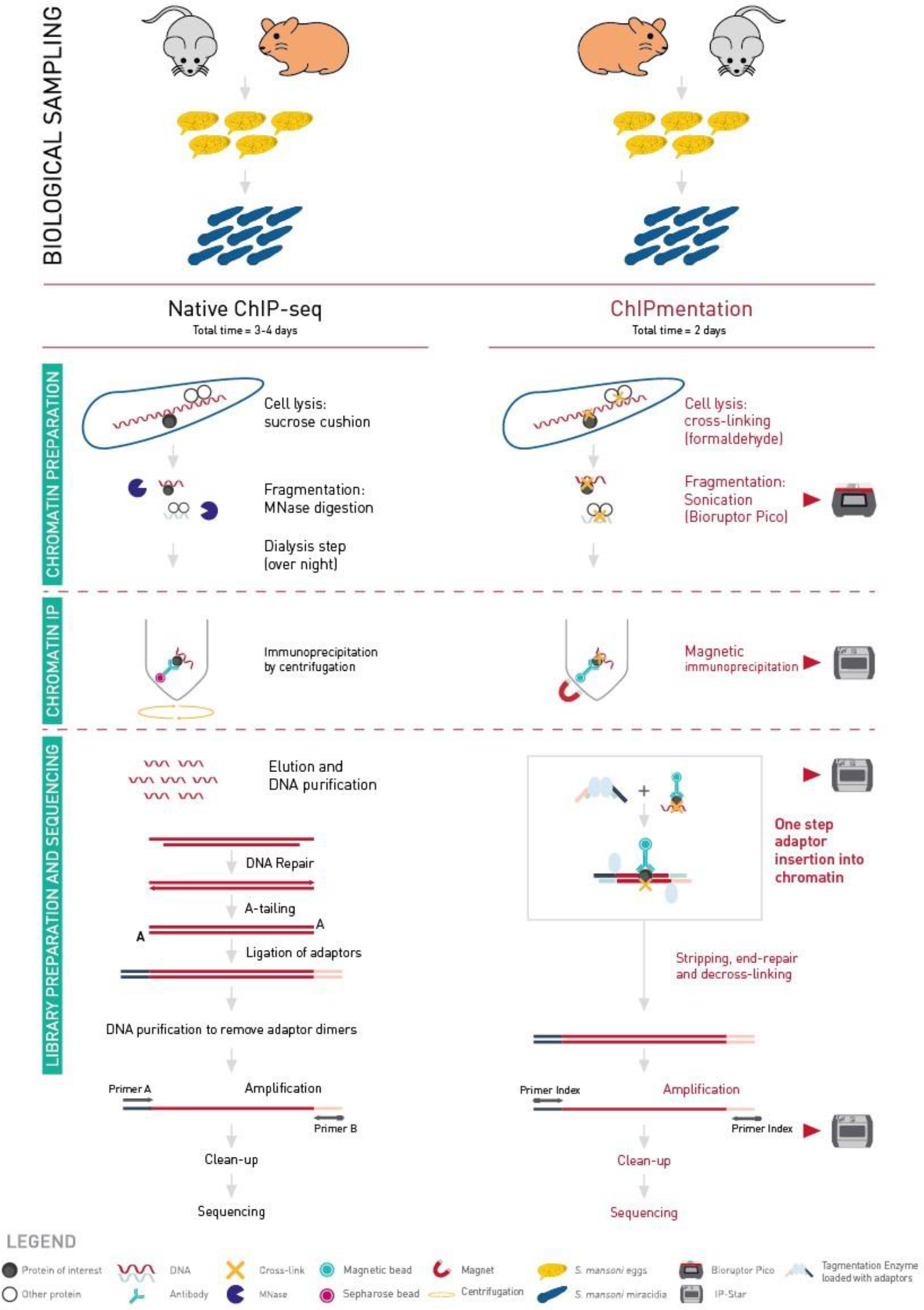
Comparison of Native-ChIP and ChIPmentation workflows. Biological sampling time depends on the biological model used. Native-ChIP protocol lasts 3 to 4 days. Sucrose cushion is used for cell lysis and MNase digestion for the fragmentation step. The immunoprecipitation is done by centrifugation. All the process is manually done. ChIPmentation protocol lasts 2 days. Cross-linking is used for cell lysis and sonication for the fragmentation step. The immunoprecipitation, tagmentation and library cleaning are done with the IP-Star.

ChIPmentation is a ChIP-sequencing (ChIP-seq) technology which uses a transposase to add the sequencing adaptors to the DNA of interest instead of the classical multi-step processing including end repair, A-tailing, adaptor ligation and size-selection (Schmidl et al. 2015). Thanks to the action of the transposase, loaded with sequencing adaptors, the library preparation is performed in only one step, which reduces hands-on time and material loss. Moreover, in the ChIPmentation approach this tagmentation process is performed directly on chromatin during the immunoprecipitation process, instead of naked DNA after purification. This workflow allows for a more reproducible tagmentation.

The combined facts that ChIPmentation has been automated on Diagenode’s IP-Star Compact Automated System and that this technology has been validated on low amounts of human cells (Schmidl et al. 2015, Roels et al. 2020) make it a perfect candidate for ChIP-seq of various targets on limited *Schistosoma mansoni* material.

## Results

### ChIPmentation has a sensitivity that is comparable to N-ChIP

10^6^-10^4^ cell equivalents gave comparable Bioanalyzer formats with peaks around 1kb, from 5,000 cells equivalent on fragments of smaller size became clearly visible. No high molecular fragments were observed in the negative control without chromatin (figure 2, table 1).

**Table 1:**
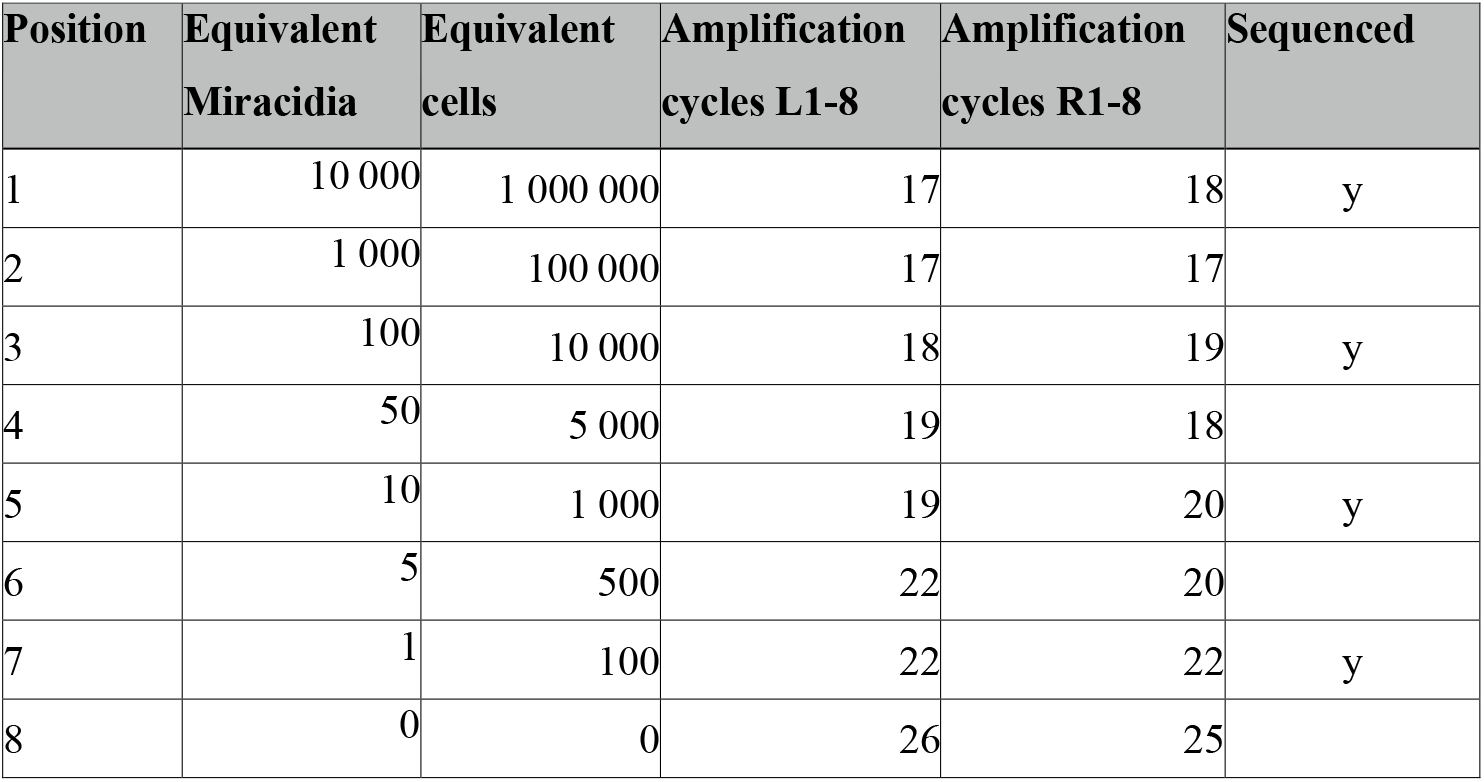
cell number equivalents and library amplification cycles for ChIPmentation libraries.

**Figure 2:**
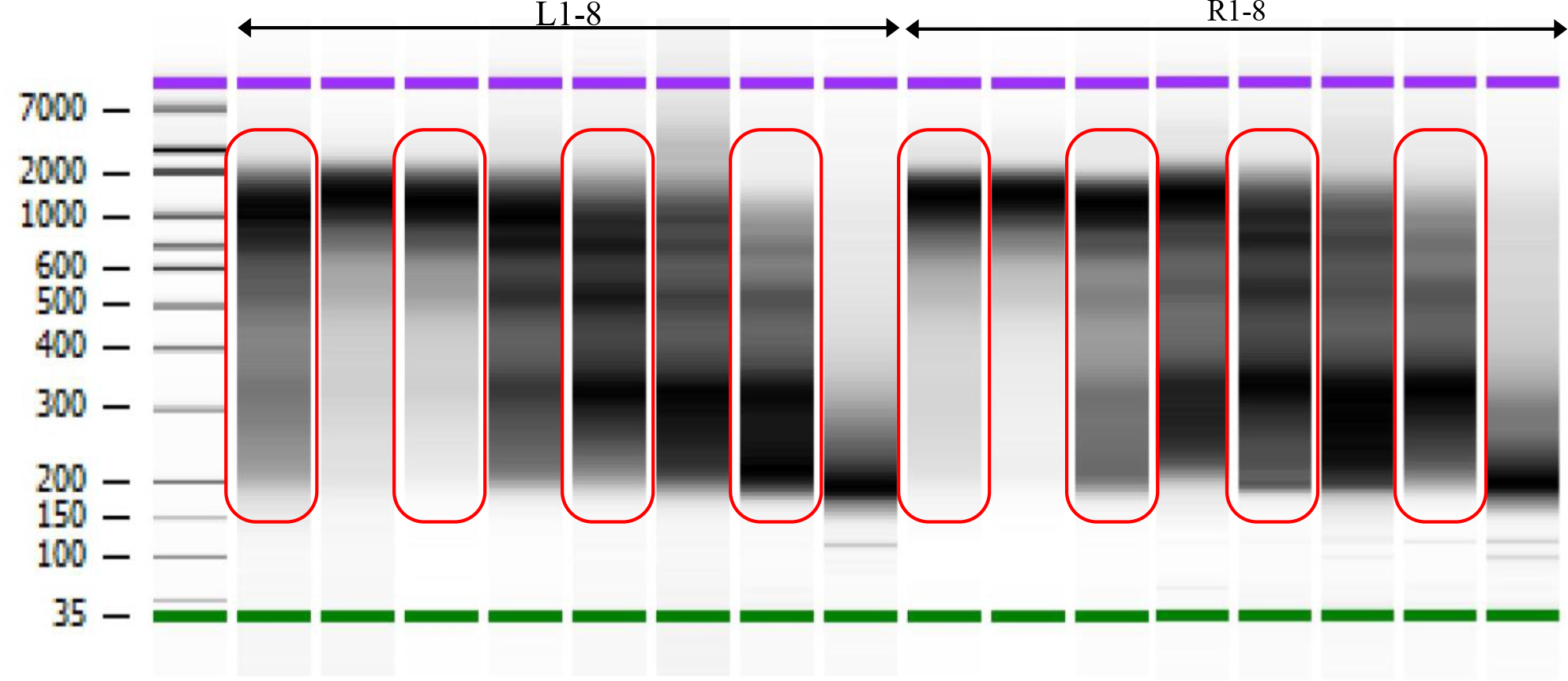
ChIPmentation library profiles for samples (L) and technical replicates of IP-Star (R). For corresponding cell numbers see table 1. Profiles surrounded in red marks those that were subsequently sequenced.

Alignment gave expected results (~50% uniquely aligned reads) for 10^6^-10^4^ cell equivalent but dropped to ~35% with 103 cells equivalent and <20% for 100 cells equivalent. No contaminating DNA was detected (table 2).

**Table 2:** General statistics on ChIPmentation libraries. All libraries were downsampled to 3.8 Mio uniquely aligned reads.

| Pos | Equivalent<br>Miracidia | Equivalent<br>cells | Amplification<br>cycles | qBit<br>DNA HS<br>ng/μl | Read pairs | Uniquely<br>aligned | After<br>deduplication | %<br>unique<br>non-dup | % to<br>keep<br>for 3.8<br>Mio |
| --- | --- | --- | --- | --- | --- | --- | --- | --- | --- |
| L1 | 10 000 | 1 000 000 | 17 | 23.2 | 5 728 173 | 7 969 894 | 6 775 525 | 59 % | 56 % |
| R1 | 10 000 | 1 000 000 | 18 | 21.2 | 8 388 553 | 13 469 770 | 10 007 402 | 60 % | 38 % |
| L3 | 100 | 10 000 | 18 | 25.2 | 12 537 243 | 19 157 585 | 14 417 614 | 57 % | 26 % |
| R3 | 100 | 10 000 | 19 | 30.4 | 8 665 837 | 13 838 684 | 8 409 793 | 49 % | 45 % |
| L5 | 10 | 1 000 | 19 | 13.8 | 11 436 716 | 17 920 411 | 8 664 467 | 38 % | 44 % |
| R5 | 10 | 1 000 | 20 | 19.3 | 11 109 380 | 17 854 360 | 7 214 153 | 32 % | 53 % |
| L7 | 1 | 100 | 22 | 22.0 | 11 662 120 | 15 737 202 | 4 095 477 | 18 % | 93 % |
| R7 | 1 | 100 | 22 | 11.9 | 10 566 906 | 15 221 913 | 3 827 034 | 18 % | 99 % |

In order to compare ChIPmentation results to Native-ChIP (N-ChIP) we re-analysed earlier data obtained by N-ChIP (Augusto, Cosseau et al. 2019) using the same data cleaning, alignment and peak calling parameters as for ChIPmentation (table 3).

**Table 3:** General statistics on N-ChIP libraries. If possible, libraries were downsampled to 3.8 Mio uniquely aligned reads.

| Pos | Equivalent<br>Miracidia | Equivalent<br>cells | Amplification<br>cycles | Read pairs | Uniquely<br>aligned | After<br>deduplication | % unique<br>non-dup | %<br>downsampled |
| --- | --- | --- | --- | --- | --- | --- | --- | --- |
| A | 8 000 | 800 000 | 14 | 13 129 021 | 11 299 195 | 10 763 378 | 41 % | 35 % |
| B | 100 | 10 000 | 14 | 21 415 343 | 5 409 960 | 2 477 749 | 6 % | 100 % |
| C | 10 | 1 000 | 14 | 29 857 139 | 12 170 093 | 6 718 570 | 11 % | 57 % |

For ChIPmentation, peakcalling with Peakranger was very robust for 10^6^ cells and delivered the expected values (based on earlier N-ChIP results). Below this cell equivalent, peak calling became dependent on bin size (table 4).

**Table 4:** Optimization of peakcalling with Peakranger. Number of peaks identified for each condition. ChIPmentation on top. 1000 bp was selected as the best extension length and applied to N-ChIP data below. N-ChIP A is for 0.8×10^6^ cells.

| cell<br>equivalents | 1 000 000 |  |  | 10 000 |  |  | 1 000 |  |  | 100 |  |
| --- | --- | --- | --- | --- | --- | --- | --- | --- | --- | --- | --- |
|  | ChIPmentation |  |  | ChIPmentation |  |  | ChIPmentation |  |  | ChIPmentation |  |
| Peakranger<br>read extension<br>length in bp | L1 | R1 | N-ChIP A | L3 | R3 | N-ChIP B | L5 | R5 | N-ChIP C | L7 | R7 |
| 100 | 6 871 | 6 767 |  | 625 | 424 |  | 200 | 370 |  | 4 732 | 5 683 |
| 300 | 10 369 | 9 545 |  | 2 125 | 1 211 |  | 499 | 635 |  | 3 944 | 5 264 |
| 600 | 10 581 | 9 924 |  | 3 513 | 2 823 |  | 610 | 813 |  | 3 197 | 3 816 |
| **1000** | **9 632** | **9 835** | **8320** | **4 033** | **2 749** | **3888** | **695** | **811** | **131** | **3 098** | **3 416** |
| 1500 | 9 901 | 9 426 |  | 3 315 | 2 925 |  | 847 | 837 |  | 4 019 | 3 526 |
| 2000 | 9 067 | 8 353 |  | 3 052 | 2 631 |  | 626 | 693 |  | 4 336 | 3 897 |

We iteratively identified 1,000 bp as best read extension length. The use of HMM-based ChromstaR improved peak calling for ChIPmentation 10^4^ cell equivalent, but not for 10^3^ or 100 cells. We obtained comparable results for ChIPmentation of 10^6^, 10^4^ cells and N-ChIP ~10^6^, 10^4^ and 10^3^ cells (table 5).

**Table 5:** Peakcalling with HMM-based ChromstaR before and after merging adjacent peaks.

| cell equivalents | 1 000 000 |  | 10 000 |  | 1 000 |  | 100 |  |
| --- | --- | --- | --- | --- | --- | --- | --- | --- |
| ChromstaR bin 1000, step 500, post prob 0.999 | all | merged | all | merged | all | merged | all | merged |
| ChIPmentation | 13 565 | **6 682** | 7 194 | **5 504** | 3 137 | **2 876** | 10 665 | 10 522 |
| N-ChIP | 6 262 | **6 186** | 7 058 | **6 922** | 5 296 | **5 129** |  |  |

ChromstaR metagene profiles showed consistent profiles for ChIPmentation 10^6^, 10^4^ and 10^3^ cells, and all N-ChIP, but not for ChIPmentation on 100 cells equivalents (figure 3).

**Figure 3:**
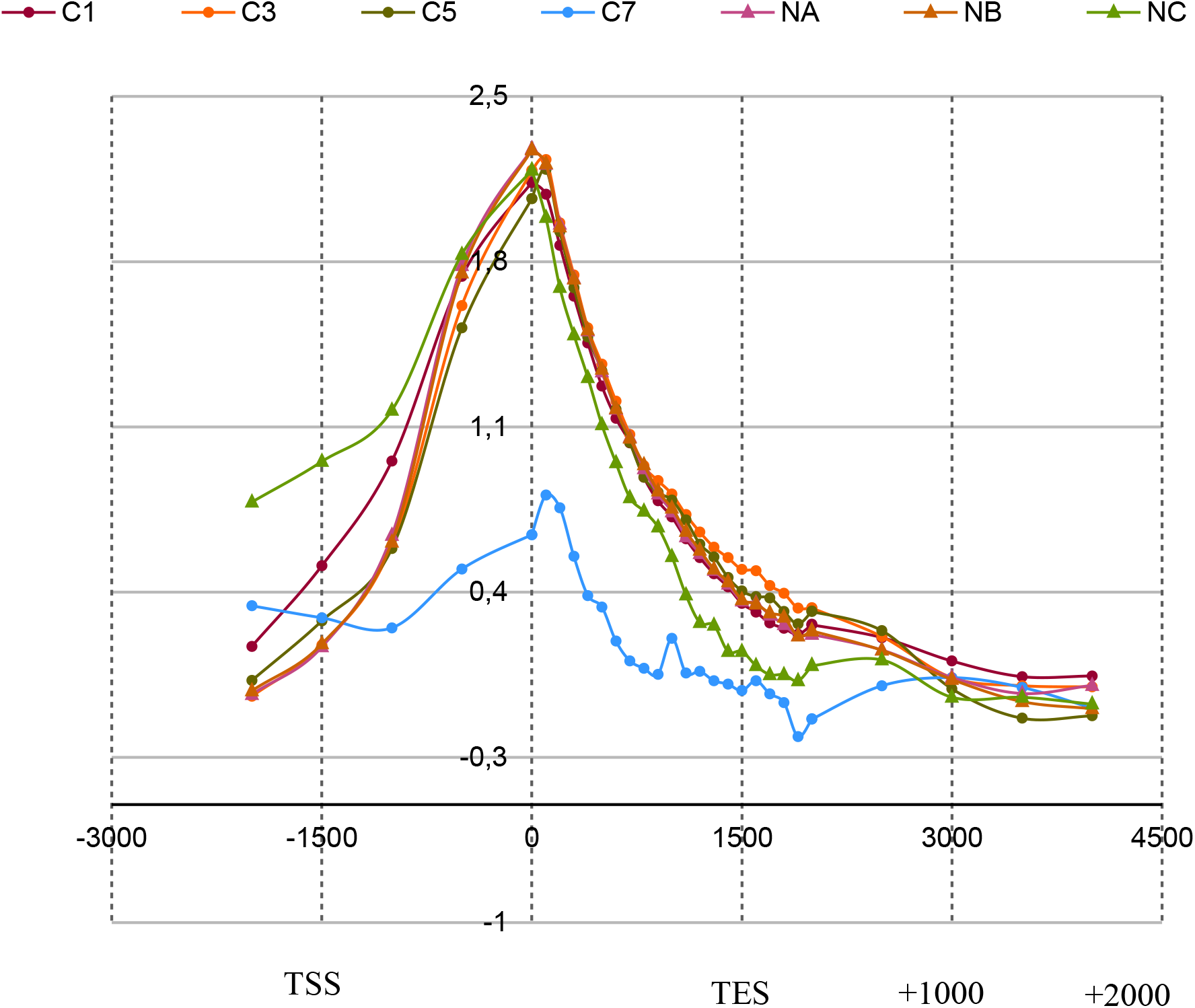
Average metagene profiles over 5,073 plus strand genes for ChIPmentation 10^6^ (C1), 10^4^ (C3), 10^3^ (C5) and 100 cell equivalent (C7), and N-ChIP 0.8×10^6^ cells (NA), 10^4^ (NB) and 10^3^ (NC). X-axis: bp upstream, within and downstream of genes. TSS/TES for transcription start and end sites. Y-axis: log(observed/expected). Not all genes contribute to the profiles as only roughly half of the genes show a H3K4me3 peak at the TSS.

ChromstaR allows for estimating correlation of chromatin profiles based on read counts (figure 4), and indicates high correlation between ChIPmentation 10^6^, and N-ChIP ~10^6^, 10^4^.

**Figure 4:**
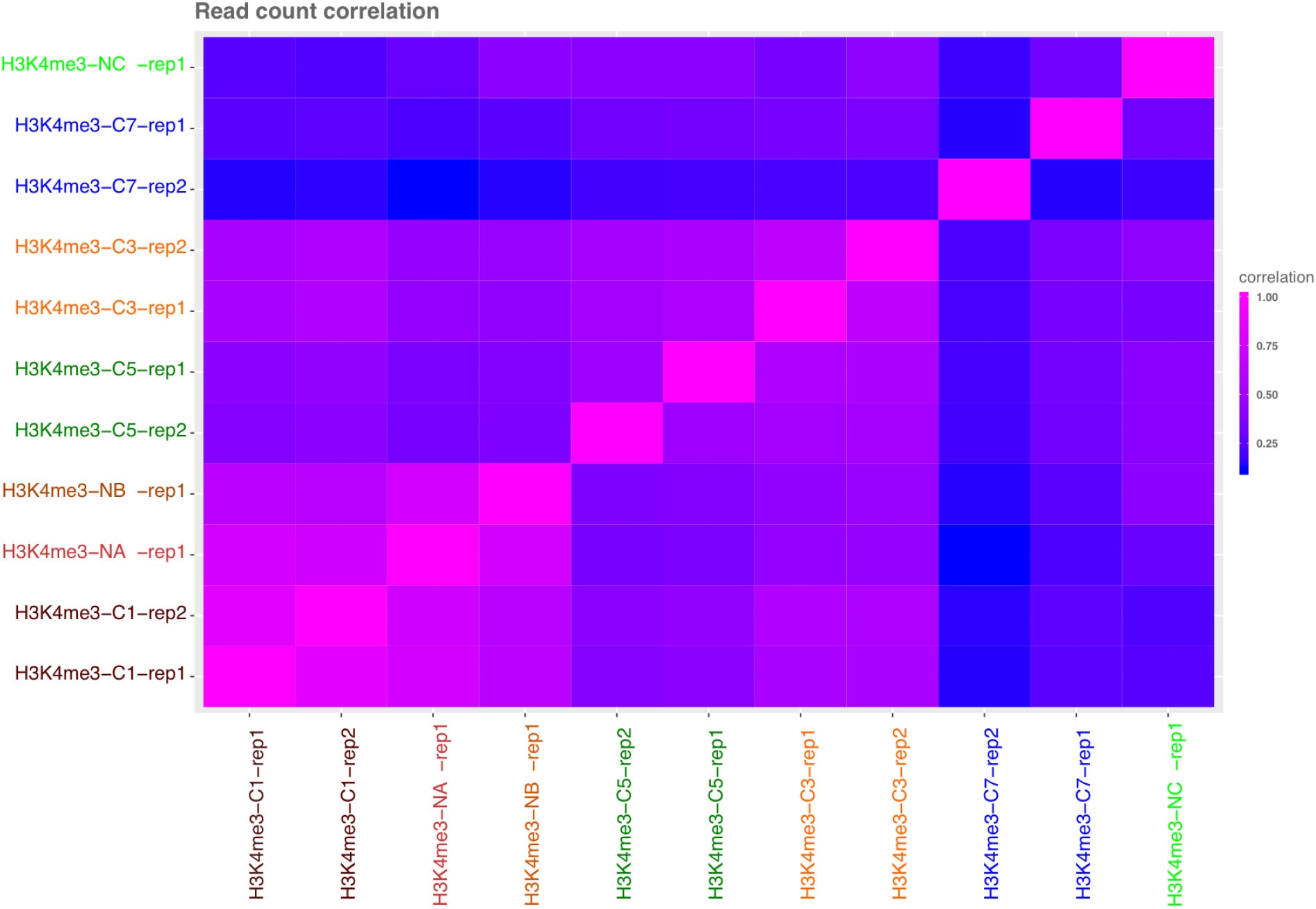
ChromstaR read count correlations between libraries (lowest 0, highest 1). L and R samples were considered as replicates 1 and 2 for ChromstaR analysis.

All other over-all chromatin profiles were below 0.75 correlation coefficient. This is surprising given the high similarity of metagene profiles. Visual inspection of peaks and profiles showed that peaks were actually correctly identified by ChromstaR (but much less by Peakranger) in ChIPmentation until 10^4^ cells but there is higher background than in N-ChIP which probably spoils correlation for lower cell equivalents (figure 5).

**Figure 5:**
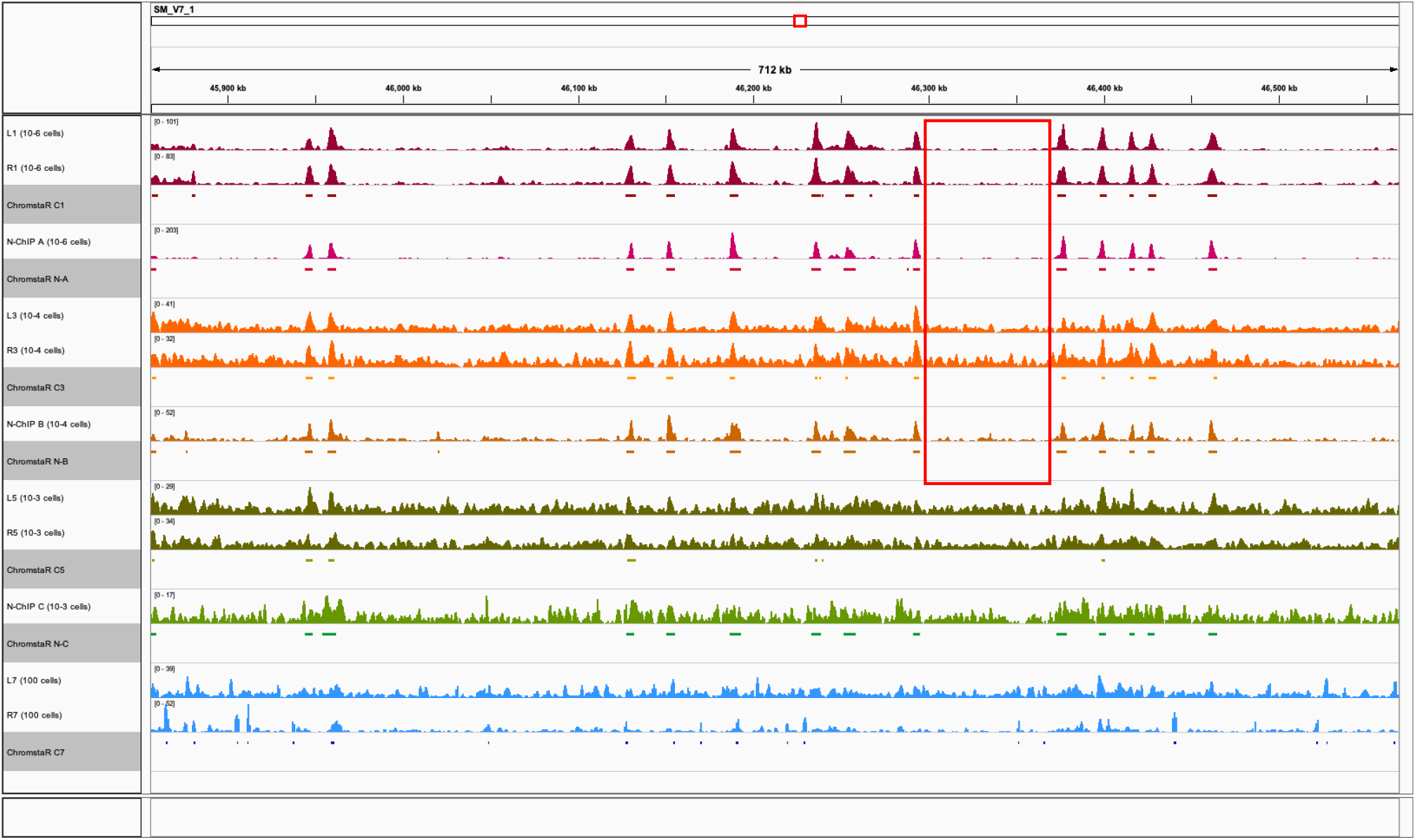
Genome browser screen shot of typical region of the *S. mansoni* genome with Peakranger chromatin profiles for visual inspection and HMM model based ChromstaR peak regions (grey underlay). ChIPmentation 10^6^ (L1, R1), 10^4^ (L3, R3), 10^3^ (L5, R5) and 100 cell equivalent (L7, R7), and N-ChIP 0.8×10^6^ cells (N-A), 10^4^ (N-B) and 10^3^ (N-C). Color codes as in previous figures. The region surrounded in red illustrates higher background for 10^4^ cells equivalent in ChIPmentation C3 (orange, replicates L3, R3) than in N-ChIP B (dark orange).

### ChIPmentation input library can rapidly be produced in parallel to the automated procedure

After having formally established that automated ChIPmentation had a comparable sensitivity to our routine N-ChIP procedure we aimed to identify the optimal way to produce input libraries for control of unspecific enrichment. The production of input chromatin is “build-in” the N-ChIP protocol (Cosseau et al. 2009, Roquis et al. 2018, de Carvalho Augusto et al. 2020) and need to be adapted to the automated ChIPmentation procedure. During a ChIPmentation experiment three types of input can be imagined (figure 6): i-1μL of chromatin before immunoprecipitation, ii-chromatin that binds non-specifically to any support and iii-available chromatin for immunoprecipitation. The aliquot of i-1μL is taken from the sample before immunoprecipitation. The two other types (input library ii and iii) need a supplementary sample in which mock immunoprecipitation is done without antibody. After immunoprecipitation, this supplementary sample contains magnetic beads with the non-specifically bound chromatin and the supernatant which is the available chromatin for immunoprecipitation.

**Figure 6:**
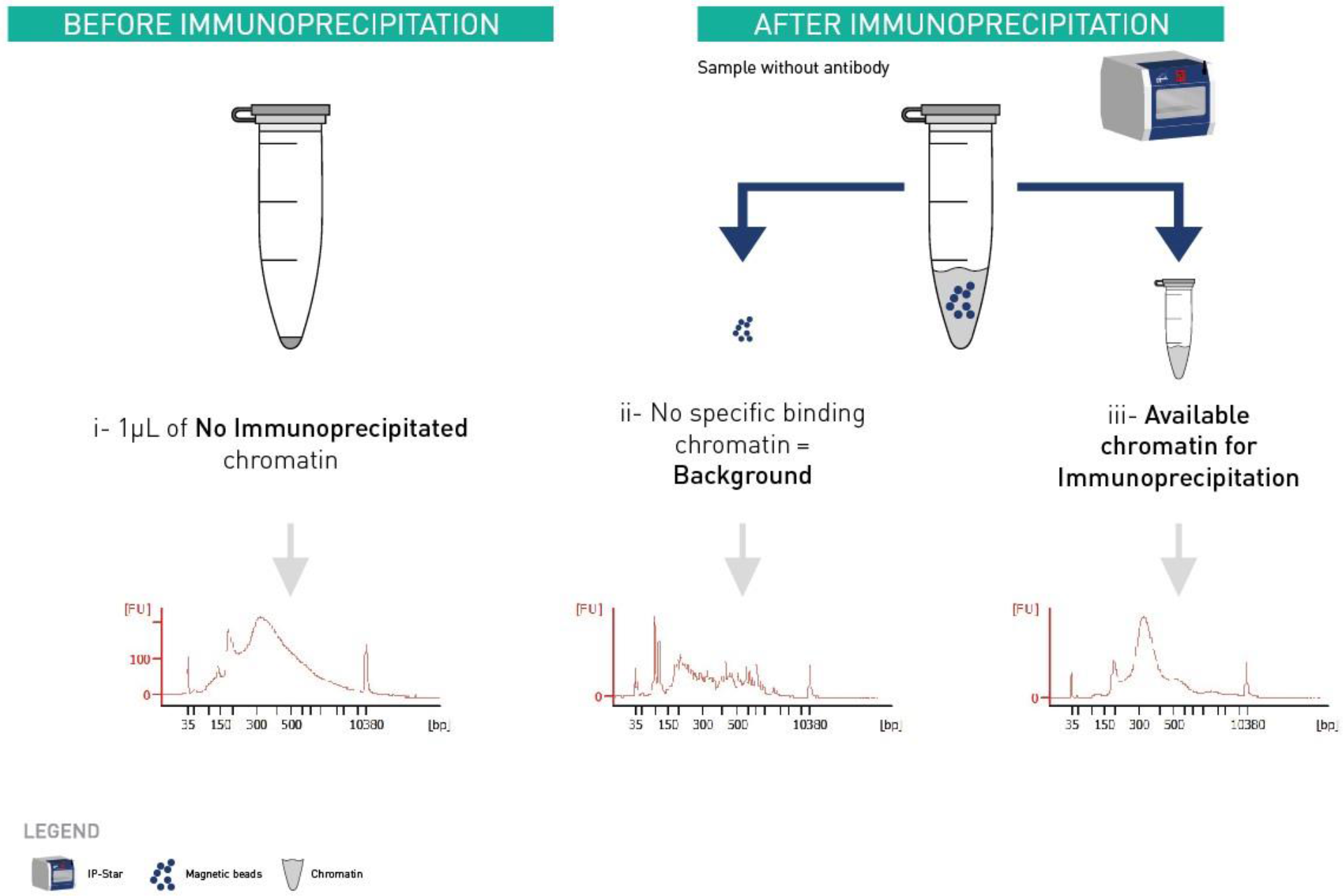
Type of inputs generated during ChIPmentation and its bioanalyser profiles associated. X axis represents the size in base pair, and Y axis represents the fluorescence intensity. Ideal library size is between 150 bp and 500 bp. i-1μL aliquot of one sample chromatin took before its immunoprecipitation. ii- and iii-are from a supplementary sample where immunoprecipitation in the IP-Star is done without antibody. After immunoprecipitation, this supplementary sample contains magnetic beads with the ii-non-specifically binding chromatin and the supernatant which is the iii-available chromatin for immunoprecipitation. ii-No specific binding chromatin gave not usable library.

Only i-before immunoprecipitation chromatin and iii-available chromatin for immunoprecipitation give ideal library sizes (figure 6). We decided to optimize the input protocol for the i-1μL of non-immunoprecipitated chromatin because it does not occupy a slot in the IP-Star. In addition, during the preliminary test, 1μL chromatin input showed a smaller Ct compare to the available chromatin input which means that it requires less amplification cycles (data not shown).

For the preparation of the 1μL chromatin input libraries, we identified two critical parameters: the first one is the dilution of the tagmentation enzyme. Using undiluted Tn5 caused complete overtagmentation. Between 10-fold and 100-fold dilutions in water delivered much better results (figure 7-A).

**Figure 7:**
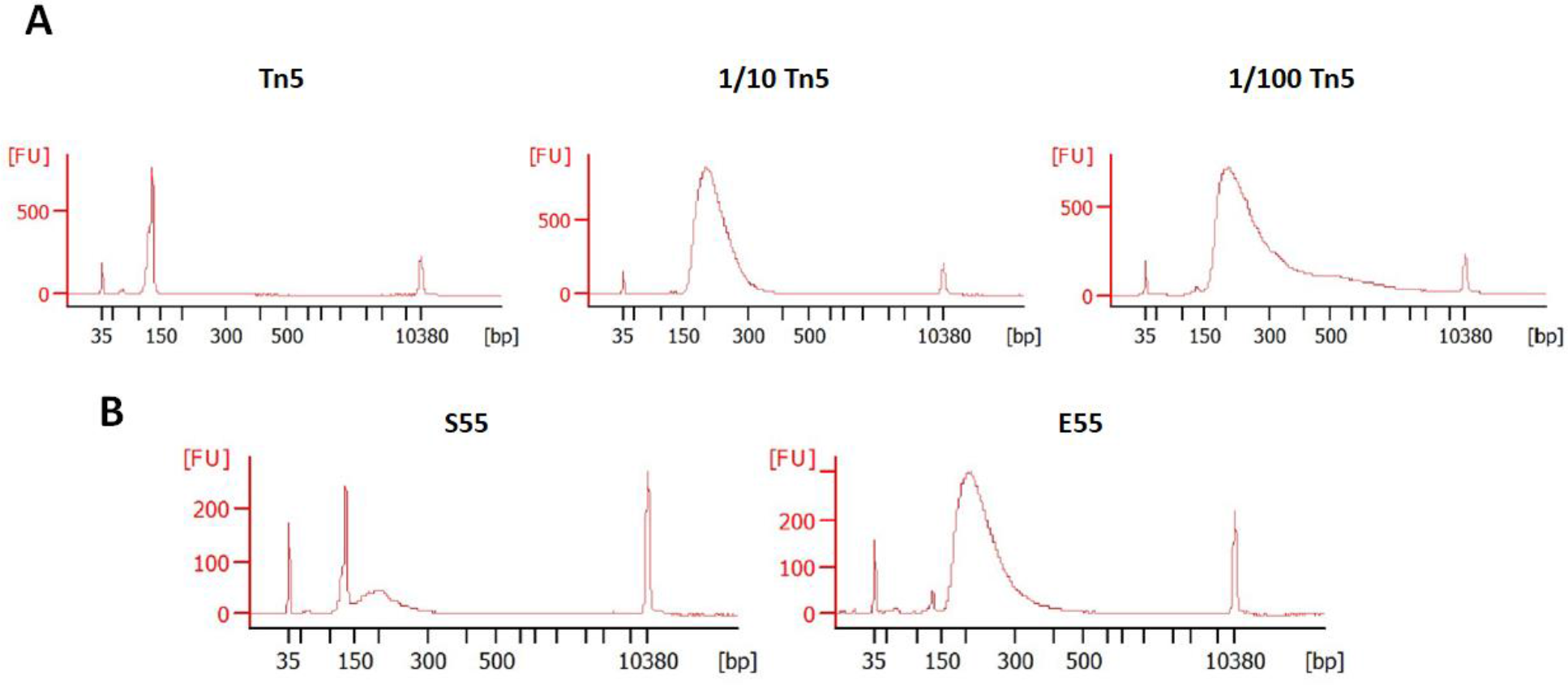
Bioanalyser profiles for input libraries done **A)** with undiluted Tn5 (**Tn5**), 10-fold diluted Tn5 (**1/10 Tn5**) and 100-fold diluted Tn5 (**1/100 Tn5**). **B)** 10-fold diluted Tn5 with 0.2% SDS (**S55**) and 10-fold diluted Tn5 without 0.2% SDS (**E55**). X axis represents the size in base pair, and Y axis represents the fluorescence intensity. Ideal library size is between 150 bp and 500 bp.

Secondly, we found that, in our hands, there is no need to add tagmentation neutralizer (0.2% SDS) after tagmentation (figure 7-B and supp figure 1). This actually inhibits the PCR amplification step (Picelli et al. 2014). Interestingly, parameters like tagmentation temperature (37°C-55°C) (suppl. figure 1), tagmentation time (2-10 min) and addition of MgCl_2_ did not have a critical effect on input libraries process.

## Discussion

Here we operationally define the limit for the robust detection of peaks with ChIPmentation to be 100,000 cells equivalent per antibody reaction, with 10,000 being the absolute limit if background is acceptable. Operational limit for N-ChIP is 10,000 cells equivalent, confirming our previous results. We therefore established ≥20,000 cells as lower limit for the routine ChIPmentation-seq procedures in *S. mansoni*. This is a bit higher than what has been described in Human cells where good results have been obtained on as low as 5,000 cells, but this is consistent with the fact that the genomic content of one cell is not the same between the two species.

To improve signal to noise ratio and reduce background in ChIPmentation it could be useful to increase the washing time (currently 10 min) and speed (currently medium) but we recommend increasing cell number than to invest into washing optimization.

1μL non-immunoprecipitated chromatin input is the best compromise to save space in the IP-Star, experiment time and biological materials when you are restricted by quantity.

This ChIPmentation protocol is not limited to miracidia cells. We also performed this protocol on *S. mansoni* adult worms and snail hepatopancreas infected by diurnal and nocturnal *S. mansoni* from Oman sporocysts (data not published). Remember that before to do any ChIPmentation experiment, it is necessary to determine the amount of antibody and the sonication time adapted to your model.

A new version of the ChIPmentation solution, called μChIPmentation for Histones (Diagenode, Cat. No. C01011011), has also been released recently in order to improve the quality on low amounts samples. This relies on a reduced number of steps, especially during chromatin preparation, and reduced number of tube transfers, in order to avoid DNA loss. It also contains a new protocol to process the non-immunoprecipitated chromatin input samples up to sequencing. This new version of μChIPmentation may be a good alternative for experiments on very low number of cells in the future.

## Material and Methods

### Production of biological material

*S. mansoni* NMRI eggs were extracted from livers of golden hamsters (gift of ParaDev) 42 days post-infection. Nocturnal *S. mansoni* from Oman (Mouahid et al. 2012) were extracted from livers of two swiss OF1 mice 8 months post-infection. Miracidia were allowed to hatch for two hours in spring water, collected by pipetting and sedimented on ice for 30 min. Miracidia were counted under the microscope, aliquoted and stored at −80°C.

### Cell lysis and chromatin shearing

Chromatin preparation was performed using the Diagenode’s ChIPmentation Kit for Histones, Cat. No. C01011009 and protocol with minor modifications. Two times 10,000 miracidia (1,000,000 cells based on the observation that 1 miracidium is composed of 100-120 cells) were resuspended each in 500 μL 1x HBSS and crashed with a plastic pestle in an Eppendorf tube on ice during ~1 min. For crosslinking, 13.5 μL of formaldehyde were added and tubes were incubated for 10 min at room temperature with occasional inversion. To stop crosslinking fixation, 57 μL of glycine were added and samples were incubated for 5 min at room temperature. Samples were centrifuged at 500xg, at 4°C, for 5 min. The pellet was resuspended in 2×1 mL of ice-cold Lysis Buffer iL1, combined and homogenized in a Dounce (pestle A) on ice for 5 min. After another centrifugation (500g, 5 min, 4°C), the supernatant was discarded and the pellet was resuspended in 1 mL of ice-cold Lysis Buffer IL2 and centrifuged (500xg, 5 min, 4°C). The supernatant was discarded and the pellet was resuspended in 100 μL of complete Shearing Buffer iS1 for each tube. Samples were sonicated with the Bioruptor Pico (Diagenode, Cat. No. B01080010) for 5 cycles (30s ON and 30s OFF). After transfer into new tubes, samples were centrifuged (16,000g, 10 min, 4°C). The supernatants were transferred into a new single tube (200 μl total) and 20 μl iS1 were added yielding 220 μl total volumes. The procedure was done in duplicates (named L and R in the following). Serial dilutions were done in iS1 to produce 100 μl equivalents of 10,000 miraciadia (106 cells), 1,000 miracidia (105 cells), 100 miracidia (104 cells), 50 miracidia (5,000 cells), 10 miracidia (10^3^ cells), 5 miracidia (500 cells) and 1 miracidium (100 cells) or 100 μl iS1 as negative control.

### Magnetic immunoprecipitation and tagmentation

IP was done on the Diagenode IP-Star Compact Automated System (Cat. No. B03000002) according to the ChIPmentation Kit for Histones User Guide and screen instructions. Antibody (Ab) coating time was set to 3 h, IP reaction to 13 h, washes to 10 min, and tagmentation to 5 min. For each sample the Ab coating mix was done with 4 μl anti-H3K4me3 (Diagenode, Cat. No. C15410003; mixture of lot A1051D and A1052D). Stripping, end repair and reverse cross-linking were done as indicated in the User Guide.

### Input library tagmentation

In the ChIPmentation Kit for Histones (Diagenode, Cat. No. C01011009) the suggested strategy is to sequence one sample immunoprecipitated with a control IgG and to use it for sequencing normalization instead of the classical input which cannot be treated exactly the same way as the immunoprecipitated samples. But IgG are negative control samples and the generation of such samples on low amounts approaches involves the use of a very high number of amplification cycles that can induce some biases. A protocol for the tagmentation of the input sample has therefore been set-up as follows.

For each immunoprecipitated sample, 1μL of sheared chromatin was kept aside before IP in the IP-Star. 1μL of MgCl_2_ (Diagenode ChIPmentation kit for Histones, Cat. No. C01011009), 8μL of molecular biology grade water, 10μL 2xTagment DNA buffer and 1μL of 100-fold in molecular-grade water diluted DNA tagmentation enzyme (Illumina 20034197, lot 20464427) were added to each 1μL input. The tagmentation reaction was performed in a thermocycler for 5 minutes at 55°C. Then, 25μL of 2xPCR NEB master mix (New England Biolabs M0541L, lot 10067165) was added to each input. The end-repair and de-cross-link were performed in a thermocycler for 5 minutes at 72°C followed by 10 minutes at 95°C. An aliquot of 2μL was taken from each input and added to 8μL of quantification mix. For each reaction, this quantification mix is composed of 0.3 μL of forward and reverse ATAC-seq primers (25μM) (suppl. table 1, Buenrostro et al. 2015), 1μL SYBR green 10X (Diagenode kit), 1.3 μL of molecular biology grade water and 5μL of 2xPCR NEB master mix. While not formally tested, leftover primers of the ChIPmentation kit could probably also be used but volume must be adjusted as the primer pairs in the kit are at 10 μM instead of 25 μM. Library amplification, purification, quality checking and sequencing steps were performed as for the immunoprecipitated samples (see below).

### Library amplification

To determine optimal number of library amplification cycles we proceeded as described in step 5.1 to 5.5 of ChIPmentation User Guide with minor modifications. We determined the number of amplification cycles for each library by using the number of cycles that corresponded to 1/3 of the slope of qPCR amplification curve during the exponential phase. Results are in table 1

Amplification was done in step 5.7. After PCR, 48 μl of library amplification mix were AMPure purified on the IP-Star using 86 μl of AMPure beads (1.8x). Instead of resuspension buffer we used ChIP grade water. After that libraries were in 20 μl.

### Library check and Sequencing

Library fragment size and concentration was checked on an Agilent 2100 Bioanalyzer with a High Sensitivity DNA Assay v1.03. Paired-end sequencing (2×75 cycles) was performed at the Bio-Environnement platform (University of Perpignan, France) on a NextSeq 550 instrument (Illumina, USA).

### Bioinformatics Analysis

Reads were quality checked with FastQC. Adapters were detected in less than 4% of reads. Reads were aligned to the *S. mansoni* v7 reference genomes using Bowtie2, sensitive settings and default values.

Uniquely aligned reads were filtered from the BAM files using the XS: tag of Bowtie. PCR duplicates were removed with samtools rmdup. BAM files were subsampled to 3.8 Mio uniquely aligned reads with picard DownsampleSam. Peakcalling was done with Peakranger P value cut off: 0.0001, FDR cut off: 0.05, Read extension length 100 - 2,000 bp, Smoothing bandwidth: 99. Delta: 0.8, Detection mode: region. ChromstaR was used with a bin size same as Peakranger extension lengths and a step size of half of bin size. Miracidia genomic DNA libraries served as input. In ChromstaR postprocessing, maximum posterior probabilities to adjust sensitivity of peak detection was set to 0.999. This keeps broad peaks intact. We reasoned that gaps between peaks that were larger than a nucleosome are not biologically meaning full and peaks were merged with BEDTOOLS when they were ≤150 bp apart (the average length of DNA in a nucleosome).

Enrichment plots over metagenes were produced over 5,073 genes on the +strand based on canonical gff v7 of *S. mansoni*.

## Data availability

Fastq files are available at the NCBI SRA Project accession number XXXX.

## Competing interest statement

Competing Interests: Diagenode is the company which developed the ChIPmentation technology. Protection Animals in Research: The laboratory has received the permit N° A 66040 for experiments on animals, from both French Ministère de l’Agriculture et de la Pêche and French Ministère de l’Education Nationale de la Recherche et de la Technologie (Décret n° 87-848 du 19 octobre 1987). The housing, breeding and care of the mice followed the ethical requirements of our country. Animal experimentation followed the guidelines of the French CNRS. The different protocols used in this study have been validated by French veterinary agency. HM possesses the official certificate for animal experimentation (N° C661101) delivered by the « Direction Départementale de la Protection des Populations » (Arrêté du 19 avril 1988).

## Acknowledgement

This study is set within the framework of the « Laboratoire d’Excellence (LabEx) » TULIP (ANR-10-LABX-41), with the support of LabEx CeMEB, an ANR « Investissements d’avenir » program (ANR-10-LABX-04-01) and the Environmental Epigenomics Core Service at IHPE.

The research was funded by the French National Agency for Research (ANR) [grant ANR-17-CE12-0005-01] CHRONOGET and CNRS.

We thank S.A.S. *ParaDev* and Julien Portela for livers of infected hamsters.

We also thank the Bio-Environment platform (University of Perpignan Via Domitia) and Jean-François Allienne for support in library preparation and sequencing

CL is a PhD candidate who receives funding from the Occitanie Region (France).

## Author contributions

Laboratory experiments: CG, CL, GM, HM, RCA

Data analysis and interpretations: ACV, CC, CG, CL, GM, HM, RCA

Manuscript redaction: ACV, AZ, CG, CL, RCA

All co-authors reviewed and edit-ed the manuscript.

## Notes

### Competing Interest Statement

Anne-Clemence Veillard and Agnieszka Zelisko-Schmidt are employees of Diagenode

